# Localization free super-resolution microbubble velocimetry using a long short-term memory neural network

**DOI:** 10.1101/2021.10.01.462404

**Authors:** Xi Chen, Matthew R. Lowerison, Zhijie Dong, Nathiya Vaithiyalingam Chandra Sekaran, Chengwu Huang, Shigao Chen, Timothy M. Fan, Daniel A. Llano, Pengfei Song

## Abstract

Ultrasound localization microscopy is a super-resolution imaging technique that exploits the unique characteristics of contrast microbubbles to side-step the fundamental trade-off between imaging resolution and penetration depth. However, the conventional reconstruction technique is confined to low microbubble concentrations to avoid localization and tracking errors. Several research groups have introduced sparsity- and deep learning-based approaches to overcome this constraint to extract useful vascular structural information from overlapping microbubble signals, but these solutions have not been demonstrated to produce blood flow velocity maps of the microcirculation. Here, we introduce Deep-SMV, a localization free super-resolution microbubble velocimetry technique, based on a long short-term memory neural network, that provides high imaging speed and robustness to high microbubble concentrations, and directly outputs blood velocity measurements at a super-resolution. Deep-SMV is trained efficiently using microbubble flow simulation on real *in vivo* vascular data and demonstrates real-time velocity map reconstruction suitable for functional vascular imaging and pulsatility mapping at super-resolution. The technique is successfully applied to a wide variety of imaging scenarios, include flow channel phantoms, chicken embryo chorioallantoic membranes, and mouse kidney, tumor, and brain imaging.

## Introduction

Ultrasound localization microscopy (ULM) is a super-resolution imaging technique that relies on the localization and tracking of intravascular microbubble (MB) contrast agents to reconstruct microvasculature. It has been demonstrated by multiple research groups that MBs can act as acoustic point sources that can be localized at a sub-diffraction precision, achieving micrometer-scale vascular fidelity at clinical frequencies without sacrificing imaging depth (1–5). Furthermore, a unique feature of ULM is the ability to measure the blood flow velocity of the microvasculature, which serves as a sensitive biomarker for numerous physiological and pathological states. The ULM MB localization protocol side-steps a fundamental trade-off between imaging resolution, which is dictated by wavelength, and imaging penetration depth, which is limited by attenuation. However, ULM introduces a new consideration between MB localization precision and acquisition time. High-fidelity MB localization requires non-overlapping MB signals; but the dilution of MBs used to achieve spatially sparse signals necessitates a long imaging duration to ensure the full perfusion of all patent vasculature (6, 7). This exasperates the pragmatic challenges associated with super-resolution vascular reconstruction, including sources of tissue motion and time-varying changes in vascular flow, which severely limit the clinical application of the technology. High concentration MB injections will perfuse through more of the vasculature over the same duration, permitting shorter imaging acquisition windows but at the expense of MB localization precision with increased spurious events and erroneously reconstructed features. Rectifying these two contradictory requirements for ULM is an area of ongoing and active research for super-resolution ultrasound.

Several ULM approaches have been specifically introduced to handle overlapping MB signals from high-concentration contrast injections. Sparse recovery strategies (8, 9), such as sparsity-based ultrasound super-resolution hemodynamic imaging (SUSHI), utilize a structurally sparse prior-representing vasculature to resolve overlapping MB point-spread functions (PSF). This localization-free approach improves imaging speed but cannot measure blood flow velocity. Compressive sensing algorithms have demonstrated the ability to resolve high density MB localizations (10, 11) and deconvolution-based approaches can iteratively shrink the imaging PSF, resulting in better localization accuracy for overlapping MB signals (12, 13). These approaches improve localization accuracy and can generate high-fidelity microvascular density maps but sacrifice the ability to track individual flowing MB trajectories. Deep-learning (DL) solutions are also gaining popularity for ULM imaging, with numerous classes of neural network architecture and training data generation showing promise in improving localization accuracy, particularly for high MB concentrations, and for accelerating the time-consuming ULM processing pipeline. Specifically, van Sloun *et al*. (14) and Liu *et al*. (15) used neural network architectures to extract point target locations representing MB centroids from B-mode images in the spatial domain. Lok *et al*. (16) and Milecki *et al*. (17) used spatiotemporal data and a fully convolutional network to improve localization confidence. These proposed DL solutions directly generate super-resolved microvessel maps but, as with the above approaches, have not been demonstrated to produce blood flow velocity maps. Thus, despite these promising results, a true high-speed super-resolution technology that can handle high concentration MB injections and offer all the features of conventional ULM (that is, microvascular structure and velocity) has not been previously reported.

Here we introduce DL-based super-resolution microvessel velocimetry (Deep-SMV), a localization free approach that provides high imaging speed and robustness to high MB concentrations, and directly outputs blood velocity measurements at a super-resolution. Deep-SMV leverages the rich spatiotemporal information embedded in ultrasound MB imaging data with a convolutional neural network including long short-term memory (LSTM) blocks to provide both structural super-resolution imaging and time-resolved velocimetry of tissue vasculature, permitting super-resolution pulsatility mapping. LSTM is a type of recurrent neural network (RNN) architecture (18, 19) which is typically characterized as neural networks with “feedback loop” mechanisms that use previous output as input and maintain internal hidden states. Compared to using feedforward networks for temporal inference, which treat the temporal dimension as an extra spatial dimension, RNNs are more suited to learning temporal features because they ensure that temporal information flows in the direction of the progression of time. By keeping internal hidden states that are constantly updated with new input, RNNs can better retain temporal information. LSTM addresses the vanishing gradient problem in classical RNNs by introducing gates to modulate information flow within the network. RNNs and LSTM have been successfully applied to various tasks such as machine translation, object tracking and motion prediction (20– 22). In this implementation, Deep-SMV was trained to recognize diffraction-limited MB signals, ranging from spatially sparse to heavily overlapping conditions, to generate super-resolved microvascular structural and velocity maps. By eliminating the need for the highly inefficient process of MB localization and tracking, Deep-SMV provides effective and fast super-resolution imaging by maximizing the spatiotemporal information from the input data. We first evaluate the performance of Deep-SMV using *in vitro* flow channel phantoms and then on *in vivo* ultrasound MB data taken from the chorioallantoic membrane (CAM) of chicken embryos, which provides an optical reference standard. We then demonstrate Deep-SMV’s ability to handle more complex microvascular dynamics by applying the technique to super-resolution velocimetry of the mouse kidney, tumor, and brain.

## Results

### Deep-SMV architecture efficiently uses spatiotemporal MB data

The LSTM-UNet structure of the Deep-SMV model (**Figure 1, Supplementary Figure 1**) utilizes LTSM blocks in the UNet bottleneck layers to take advantage of the spatiotemporal information present in the input ultrasound data. The feature extraction path of the conventional UNet architecture functions to generate spatial feature maps from each frame of the input sequence without awareness of their temporal relations. The LSTM units (**Supplementary Figure 1, top**) perform spatiotemporal inference on the sequence of temporally contiguous features, generating feature maps consisting of both spatial and temporal information. Finally, the reconstruction block of the UNet performs predictions based on the spatiotemporal feature output by the LSTM units. This approach allows for the Deep-SMV network to directly train for the output of blood flow velocity from the raw spatiotemporal features, without explicitly relying on inefficient MB localization and tracking processes. This permits the network to use high concentration and overlapping MB signals more efficiently for large vascular lumens while retaining high fidelity for microvascular reconstruction.

**Figure 1.**
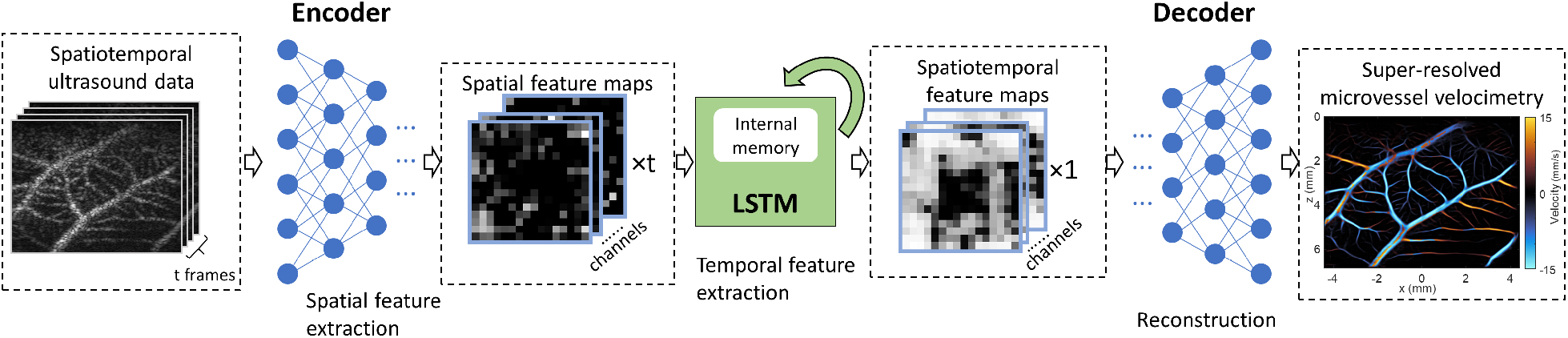
Deep-SMV data workflow. The Deep-SMV network takes a spatiotemporal (2D spatial + time) input of ultrasound data, passes this through encoder blocks to extract spatial features, uses LSTM blocks to extract temporal features, and then reconstructs a super-resolved velocimetry map from the spatiotemporal features via decoder blocks. Further details are presented in **Supplementary Figure 1**.

An example of *in vivo* informed data simulation for Deep-SMV training is demonstrated in **Supplementary Figure 2**. CAM optical images (**Supplementary Figure 2a**) provide a physiologically relevant *in vivo* vascular structure that can be readily segmented (**Supplementary Figure 2b**) to generate binarized representations of the vessel space. The process of medial axis skeletonization can then yield structural connectivity maps (**Supplementary Figure 2c**) as well as distance transform maps (**Supplementary Figure 2d**) of this vessel bed, which is informative of the local vascular diameter. This serves as the basis for an undirected vessel graph for MB flow simulation (**Supplementary Figure 2e**), which is assigned direction using a breadth-first search (BFS) traversal algorithm (**Supplementary Figure 2f**). The resulting MB locations were fed into a Field-II (23, 24) simulation (see **Supplementary Figure 2g**, detailed in **Methods**) to generate training samples of 16 frames in the temporal dimension, and 256 * 256 pixels in the spatial dimension. The loss of the NN on the training set and the validation set gradually decreases as a function of time, showing no sign of overfitting until around 75 epochs, when the validation loss plateaus, and training loss continues to show minor improvement. The trained network was tested on 32 simulation samples of 16 * 256 * 256 simulation data. It was able to achieve a mean absolute prediction error 1.59 mm/s for flow velocity prediction.

### Flow channel phantom validates Deep-SMV reconstructions

We then tested the performance of Deep-SMV on an *in vitro* flow channel phantom (**Supplementary Figure 3**) where the ground-truth velocity is known from the flow volume rate of the connected syringe pump. A contrast power image of the flow channel (**Supplementary Figure 3a**) demonstrates that the flow channel was well perfused with MB contrast agent, yielding a high MB concentration scenario where conventional ULM does not perform efficiently. This is apparent in the conventional ULM velocity reconstructions (**Supplementary Figure 3b**) where MB localization and tracking results in relatively sparse tracing of the flow channel lumen. Deep-SMV was also used to process the same dataset (**Supplementary Figure 3c**). Deep-SMV demonstrated a better realization of the parabolic flow profile within the flow channel phantom with a much higher perfusion efficiency. In comparison to the mass-conservation calculated ground-truth (**Supplementary Figure 3d**), conventional ULM showed an underestimation bias for velocity estimates, particularly for high flow volume rates, whereas Deep-SMV was more consistent across all tested flow velocities. The analysis regions of interest (ROIs) were placed to avoid the upper edge enhancement artifact from buoyant MBs at low flow volume rates (arrow). Each 1,600-frame flow volume dataset took approximately 442 seconds to process using conventional ULM without MB separation (25), and only 190 seconds to process using Deep-SMV. These processing time benchmarks include SVD filtering and data transfer times.

### Deep-SMV outperforms conventional ULM on short data segments

The use of Deep-SMV for the generation of super-resolved velocity maps takes advantage of the fast forward-processing speed of NNs and GPU-based parallelized computing to achieve real-time super-resolution imaging capability. The example 300 × 300-pixel image patch with 64 temporal frames in **Figure 2a** took approximately 2,500 ms to undergo the conventional ULM localization and tracking steps (**Figure 2b**) to produce the sparsely populated super-resolution flow velocity map shown in **Figure 2c**. The direct temporal accumulation of MB signals, a conventional diffraction-limited ultrasound technique shown in **Figure 2d**, demonstrates that most of the vascular luminal space is missed in conventional ULM for a short data segment due to the inefficient MB localization process. Applying Deep-SMV to the same data segment yielded a super-resolution flow velocity map (**Figure 2e**) which only took 28 ms to process on 2 NVIDIA® GeForce® RTX 2080 Ti GPUs, a near 100X acceleration compared to conventional ULM. The magnitude of flow estimated via Deep-SMV is comparable to the point estimates derived from ULM; however, it is notable that the proportion of vascular lumen space that was reconstructed is much higher than conventional processing. The convention used in all results presented is that the orange color indicates flow toward the transducer, and blue indicates flow away from the transducer.

**Figure 2.**
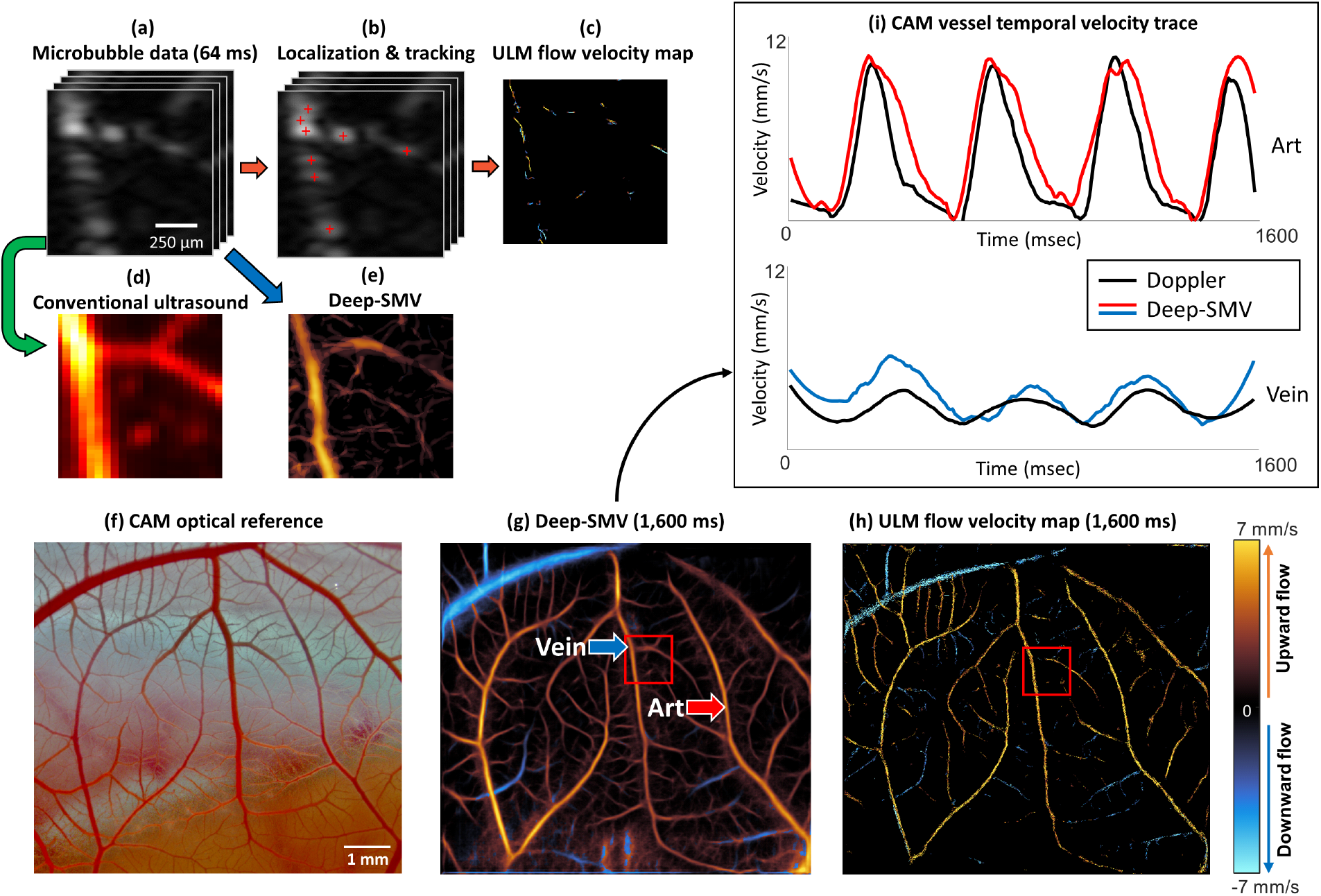
Deep-SMV workflow on CAM surface vessel data. **(a)** An example short data segment of MB data (64 ms) was processed using **(b)** conventional ULM localization and tracking, resulting in **(c)** a very sparse flow velocity map where it is difficult to visualize the microvessel anatomy. **(d)** Direct accumulation of MB signal data reveals diffraction-limited vessel structure. **(e)** Processing the same data segment with Deep-SMV allowed for super-resolution velocity estimation with more structural connectivity than conventional ULM. The orange color in (c) and (e) indicates flow toward the transducer, and blue is flow away from the transducer. **(f)-(h)** Deep-SMV and ULM performance comparison on long data segments with large FOV. **(i)** Deep-SMV shows distinct pulsatility features in local regions sampled from feeding (**Art**) and draining (**Vein**) vasculature, which matched conventional Doppler velocity measurements.

### Chicken embryo CAM velocimetry reveals super-resolved vascular pulsatility

After demonstrating good performance in the flow phantom experiment (**Supplementary Figure 3**) and for small *in vivo* imaging patches (**Figure 2a-e**), we then sought to test Deep-SMV on longer duration *in vivo* contrast ultrasound data from a large imaging field of view (FOV). We have previously shown that the CAM is an excellent model for testing ultrasound super-resolution processing methods (25–27), providing low attenuation and minimal tissue motion, while also being accessible to optical imaging to serve as a reference standard (**Figure 2f**). Deep-SMV processing of a 1600-frame (1,600 ms) IQ data acquisition (**Figure 2g**) reveals an interdigitated vessel structure that is characteristic of the CAM vascular topology, and which shows good correspondence to the optical reference. The small branching arterioles on the order of 20-30 μm in diameter, which are adjacent to the main site of gas exchange, are visible in the Deep-SMV reconstruction of this planar vascular membrane. The conventional ULM reconstruction of this same dataset (**Figure 2h**) shows comparable performance for larger vessels but was less able to distinguish the smaller interdigitated vasculature on the CAM than Deep-SMV due to the relatively short data acquisition time. The ULM processing time was approximately 1,800 seconds for this 1,600-frame acquisition, including the application of MB separation filters, SVD filters, MB pairing and tracking, and data accumulation. In comparison, the Deep-SMV image of this CAM network, which was generated by accumulating multiple 16ms subsets of data for a total duration 1.6 seconds, took only 294 seconds to process. In addition, Deep-SMV had the added advantage of providing a higher temporal resolution. As demonstrated in **Figure 2i**, Deep-SMV revealed a cyclic pulsatility in the blood flow velocity within two selected vessel regions (labelled **Art** and **Vein**) that were not apparent in conventional ULM processing. The frequency of the change in blood velocity was around 2-3 Hz, which matches with the Doppler fluctuation, and is close to the reported values for the cardiac cycle of chicken embryos at this stage of development (28).

To further investigate the utility of Deep-SMV for super-resolved pulsatility mapping, we performed a detailed CAM vessel flow dynamics analysis in **Figure 3. Figure 3a** demonstrates selected imaging frames (160 ms between each frame) from a Deep-SMV pulsatility video (**Supplementary Video 1**). At the beginning of the cardiac cycle in the chicken embryo, the main feeding vessels to this vascular bed accelerate to a peak velocity and demonstrate a rapid cyclic shift in the Deep-SMV velocity estimates over time. In contrast, the draining vessels show a much slower peak velocity, with less dramatic cycling and a slight phase delay in peak velocity time. **Figure 3b** demonstrates a local analysis performed on two regions within the accumulated Deep-SMV velocity map. In **Region 1**, we note that there is a gradual decrease in the peak velocity and pulsatility in the velocity profiles (**Figure 3c**) that were generated along the vessel length (**Figure 3c i-iv**). **Region 2** demonstrates the difference in peak velocity for the vessel cross-sections labeled **Art** and **Vein**, with a phase delay in the peak velocity estimate time apparent in the heat maps demonstrated, implying that MB flow needed time to cross the capillary space between these two vessels.

**Figure 3.**
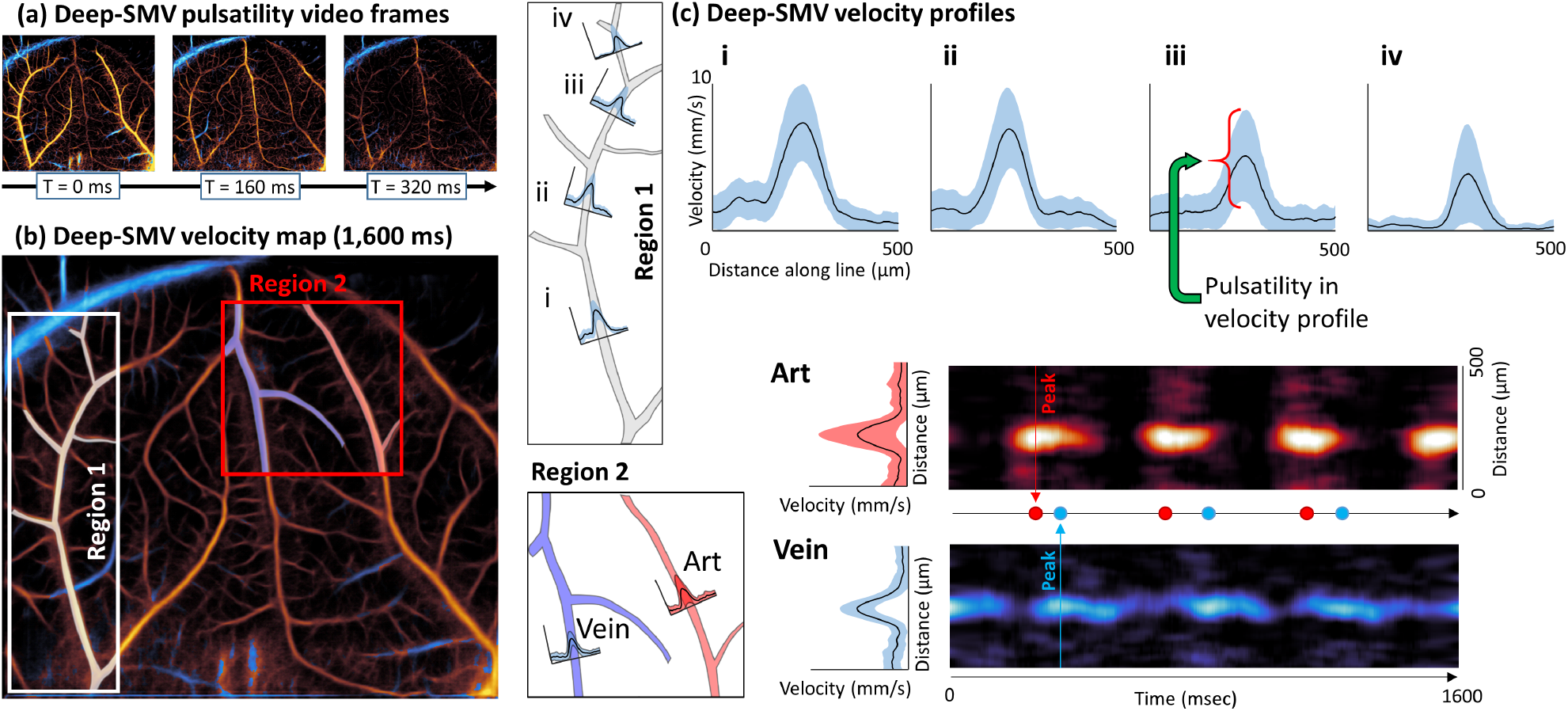
Detailed CAM flow dynamics analysis using Deep-SMV velocity estimates. **(a)** Video frames taken from a Deep-SMV pulsatility video (**Supplementary Video 1**) which demonstrate cycling pulsatility and different flow dynamics for feeding and draining vessels. **(b)** Accumulated Deep-SMV velocity map with two selected regions. Region 1 demonstrates a gradual decrease in **(c)** peak velocity and pulsatility in velocity profiles along the vessel length (**i** to **iv**). Region 2 demonstrates a phase delay in the peak velocity estimate for vessels labeled **Art** and **Vein**.

### Tissue-specific training data improves the performance of Deep-SMV

Next, we applied Deep-SMV to the extremely complex vascular structure that is present in the mouse brain. The densely packed and hierarchical vascular organization of the mouse brain presents a substantial challenge for super-resolution vascular imaging, typically requiring very low concentration MB injections and prohibitively long data accumulation times to accurately reconstruct cerebral features. A case point for this difficulty is evident in the example shown in **Figure 4a**, a short duration (1,600 ms) ULM reconstruction of the brain vasculature. This example exemplifies the graduality of ULM accumulation, with the sparsity of the reconstruction evident in both the columnar cortical vessels and the subcortical supply vasculature. Only after a substantially longer accumulation time does mouse brain hierarchical vasculature structure become apparent in conventional ULM reconstruction (**Figure 4b** – requiring 64,000 ms of data). Sporadic MB localization errors are also evident, particularly in the largest vessels, leading to spurious velocity estimates which bias the final velocity map. In comparison, the Deep-SMV network used in the previous section was used to reconstruct a velocity map using only 1,600 ms of data (**Figure 4c**), which demonstrates multiple scales of vasculature, with obvious branching connectivity ranging from the interior feeding arterioles to the cortical capillary layers – highlighting Deep-SMV’s more efficient use of high concentration MB signal.

**Figure 4.**
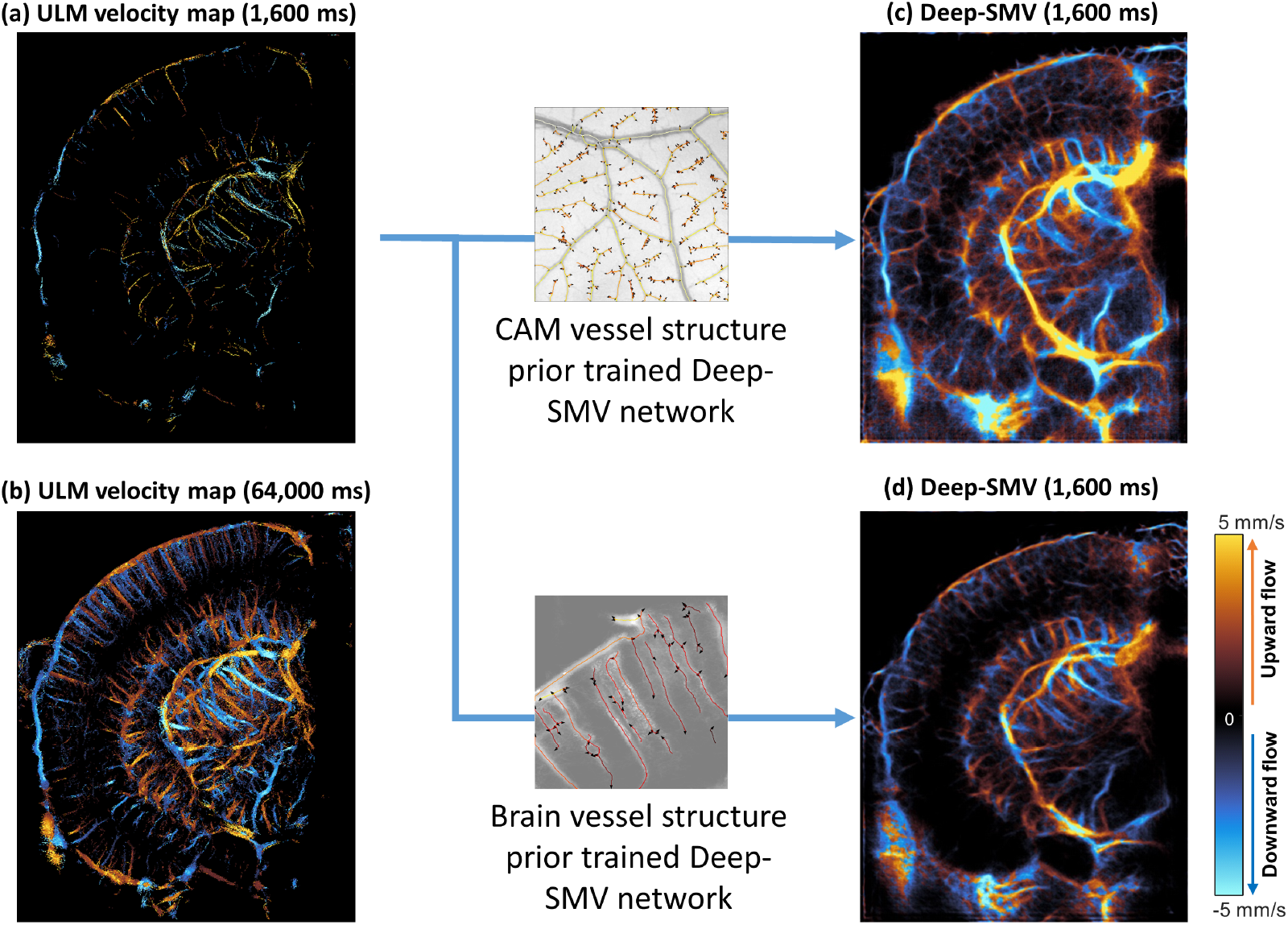
Deep-SMV with different vessel structure prior. **(a)** Conventional ULM localization and tracking for a 1600ms mouse brain dataset was only able to sparsely populate the image with velocity estimates. **(b)** Due to the inefficient use of MB signals, a much longer acquisition time (64,000ms) was required to reconstruct the majority of brain vasculature. **(c)** Deep-SMV trained with CAM simulation data had acceptable performance for larger subcortical vessels but was unable to separate parallel cortical vessels. **(d)** The mouse-brain prior trained Deep-SMV network had improved performance in the cortical regions, while maintaining the ability to reconstruct subcortical features.

In order to study the effect of a structural prior in the training data, we re-trained the model from scratch using a mouse brain ULM based simulation training set and applied it to mouse brain data. While the CAM-trained network has acceptable performance on larger vessels (**Figure 4c**), it was unable to separate parallel cortical vessels because such structure is rarely seen in the CAM template. Using mouse brain ULM trained Deep-SMV model, most of the cortical vessels can be resolved with improved clarity (**Figure 4d**). We thus concluded that the performance of Deep-SMV is indeed affected by the structural bias in the training data. However, we note that in cases where a vascular model of the organ of interest cannot be easily obtained, using training data from a different organ is still a feasible option as long as the scales of the vasculature are comparable.

### Mouse brain Deep-SMV pulsatility follows cardiac cycle

Motivated by Deep-SMV’s ability to visualize super-resolved vascular pulsatility in the CAM (**Figure 3**), we applied a similar flow dynamics analysis to a Deep-SMV reconstruction of mouse brain vasculature (**Figure 5**). Velocity line profiles taken from cortical arterioles (**Figure 5a**) demonstrated a well-developed laminar profile with cyclic pulsatility. Likewise, velocity line profiles could also be generated for deep sub-cortical vessels, such as in and around the thalamus. Segmentation ROIs were placed around cortical and sub-cortical vessels (**Figure 5b**) to generate velocity traces over the course of the 1,600 ms imaging acquisition. The pulsatility of the Deep-SMV velocity estimates (**Figure 5c**) matched electrocardiogram (ECG) measurements taken from this mouse. In the cortex, the ECG r-wave peak led the peak velocity measurement by an average (± standard deviation) of 71.1 ± 15.1 ms. For the subcortical regions, the upward flowing vessel reached peak velocity 65.5 ± 18.7 ms after the ECG r-wave peaks, and the downward flowing vessel had the largest delay in peak velocity at 90.3 ± 29.6 ms.

**Figure 5.**
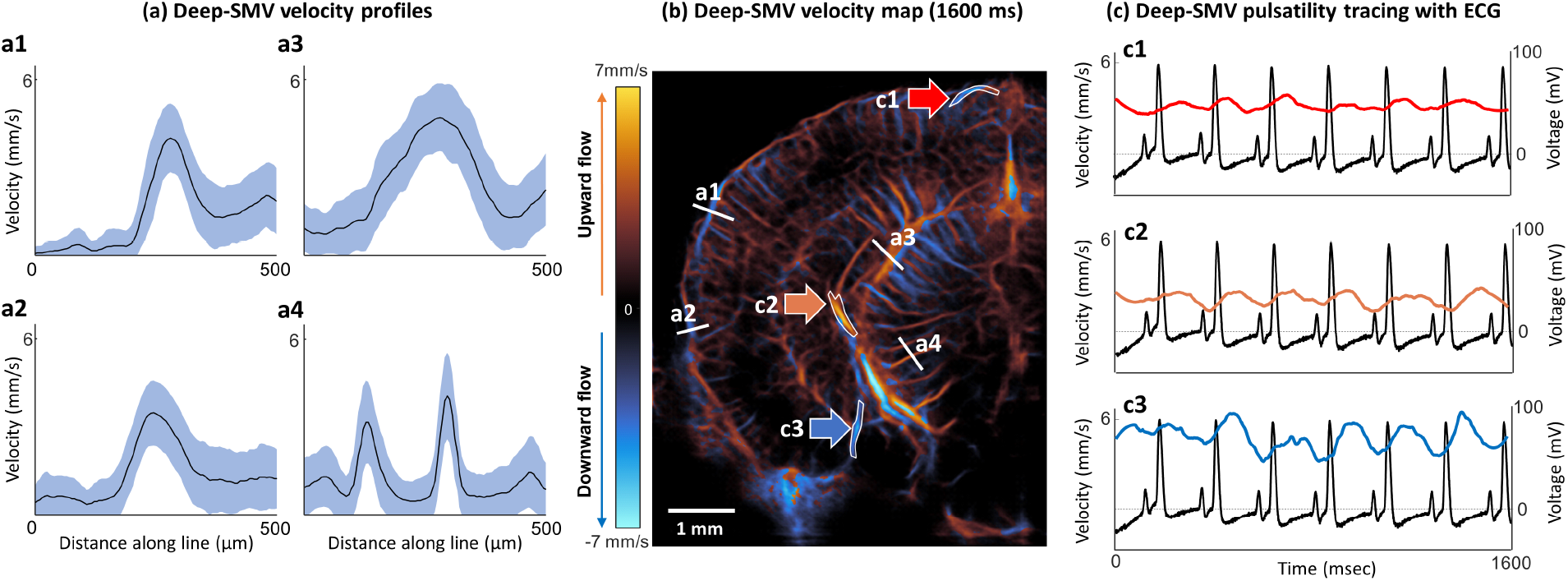
Mouse brain vessel velocimetry validated with ECG. **(a)** Velocity profiles generated from cortical (**a1** and **a2**) and subcortical (**a3** and **a4**) vasculature, taken from **(b)** a Deep-SMV velocimetry map of a mouse brain. **(c)** Velocity traces of cortical vessel (**c1**) and subcortical vessel (**c2** and **c3**) segmentations in comparison to ECG measurements.

### Deep-SMV demonstrates hierarchical in vivo vasculature in diverse organ types

Finally, we applied CAM-trained Deep-SMV to other complex *in vivo* vascular structures, namely the mouse kidney and a murine tumor xenograft model, to test the generalizability of super-resolution velocimetry performance. Contrast power Doppler of the mouse kidney (**Figure 6a**) demonstrates a typical renal anatomical vascular feature of large, deep feeding vessels that branch outward into the medulla as a fine, densely packed microvascular bed approaching the renal cortex. Applying color Doppler processing to this dataset (**Figure 6b**) reveals the directionality of the blood flow but lacks fidelity for the smallest vessels evident on the power Doppler image. Conventional ULM processing was able to capitalize on the relatively low MB concentration in this dataset to reconstruct microvessel velocity flow information (**Figure 6c**) but did not perform as well for the larger vessels in the FOV where the local MB concentration was higher. It took approximately 7,200 seconds of processing time to apply the MB separation filters, localize and track MBs, this 900-frame dataset (respiratory gating was applied to remove high tissue motion from three 720-frame acquisitions). In comparison, Deep-SMV only took 135 seconds to generate the image presented in **Figure 6d**. The Deep-SMV image also demonstrated super-resolved velocity estimation with a higher fidelity than color Doppler imaging, while outperforming ULM in the larger vessels of the mouse kidney. Deep-SMV was able to better handle the overlapping MB signals in the large renal vessels to fill in the vascular luminal space more appropriately. Similar findings are presented for a mouse Hepa1-6 tumor in **Figure 6**. Contrast power Doppler demonstrated high peripheral vasculature (**Figure 6e**) and color Doppler processing was only able to detect the larger vessels (**Figure 6f**). Conventional ULM reconstructed a very sparse vascular bed (**Figure 6g**) using a dataset of 1,600 frames and took over 9100 seconds to process. In comparison, Deep-SMV produced a superior visualization of larger vessels and could better depict the smaller tortuous microvessels in this tumor (**Figure 6h**). Deep-SMV processing took approximately 580 seconds to accumulate this image.

**Figure 6.**
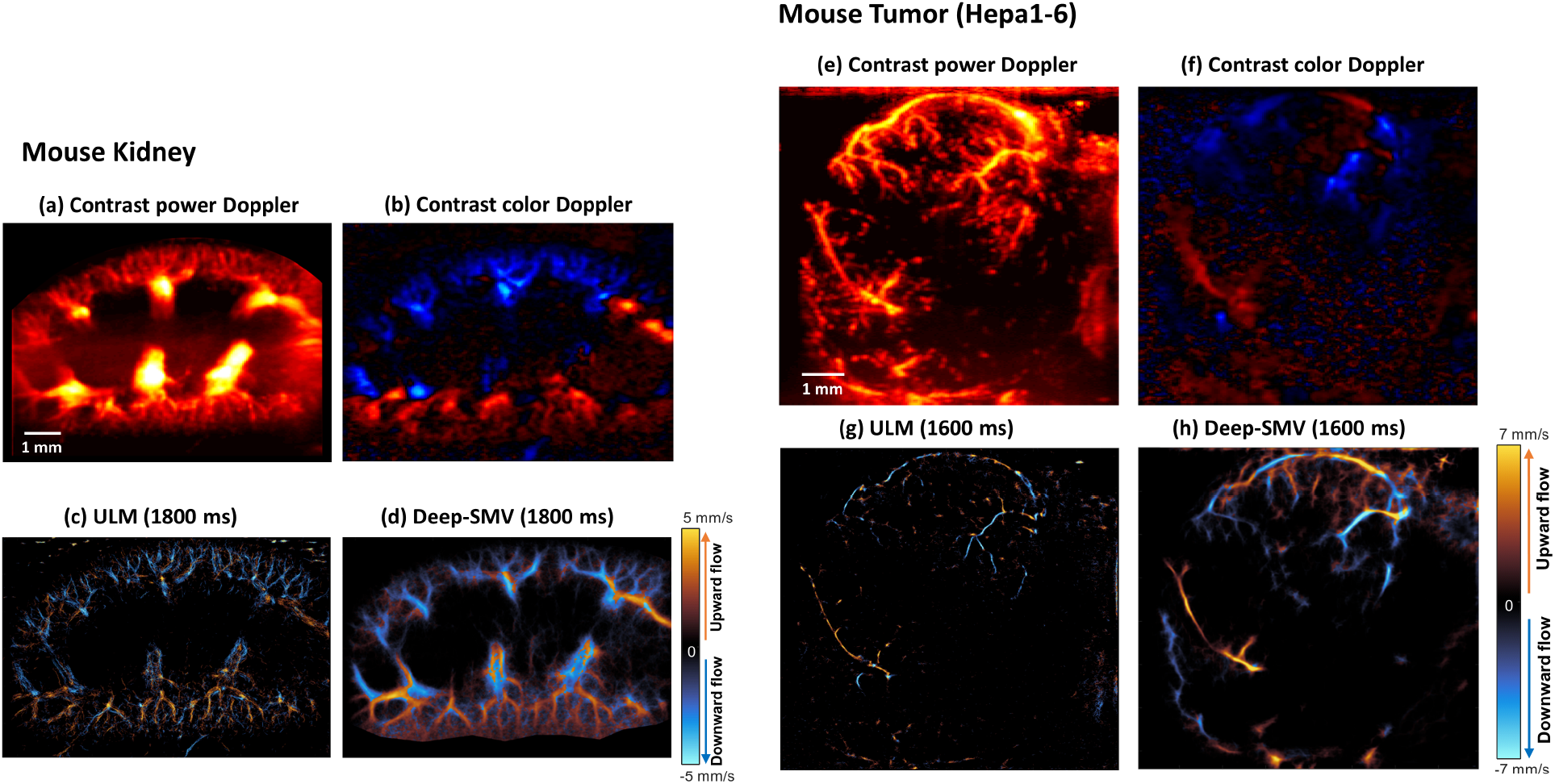
Mouse kidney and mouse tumor results. **(a)** Contrast power Doppler imaging of the mouse kidney. **(b)** Color flow Doppler demonstrates the directional flow of the larger renal arterioles and venules. **(c)** Conventional ULM imaging was able to detect some of the smaller vessels in this imaging cross section, with relatively poor performance in the larger vessels. **(d)** Deep-SMV demonstrates directional velocity of both the large and small vessels, with better visualization of the luminal space of the renal arteries. **(e)** Hepa1-6 tumors demonstrated relatively high vessel density in the periphery with reduced intratumoral vascularization. **(f)** Color Doppler could only detect the larger vessels, performing poorly in the deeper regions of the tumor. **(g)** ULM imaging of this acquisition demonstrates relatively sparse vascular reconstruction in comparison to **(h)** Deep-SMV processing which had superior visualization of larger vessels and could demonstrate better connectedness in the smaller tortuous microvessels.

## Discussion

In this study, we introduced a DL-based super-resolution microvessel velocimetry approach for contrast-enhanced ultrasound imaging, named Deep-SMV, by employing LSTM blocks in the bottleneck layers of a UNet architecture to take advantage of the rich spatiotemporal information that is present in ultrafast ultrasound MB data. Using this architecture, we can bypass the extremely expensive MB localization and tracking processing in conventional ULM to provide super-resolved blood velocity measurements that are more robust to high MB concentrations and at a high imaging speed. The training data for the Deep-SMV network was generated using ultrasound MB flow simulation on *in vivo* optical images taken from the CAM of chicken embryos and mouse brain ULM images. The technique was validated using a flow channel phantom, where less velocity estimation bias in comparison to ULM was noted, and we demonstrated that Deep-SMV can produce super-resolved velocity maps for several challenging *in vivo* imaging scenarios, including chicken embryo CAM imaging, and mouse brain imaging, mouse kidney imaging, mouse tumor imaging.

Previous research groups have applied CNNs to the problem of extracting useable ULM localization data from overlapping MB signals (14–17); however, none of these approaches yield velocity information. While it is possible to perform inference on spatiotemporal data with a conventional CNN by treating the temporal dimension as multiple channels or as an additional spatial dimension, the LSTM-based spatiotemporal model has several advantages over conventional CNN approaches. The scale of temporal perception of a multi-channel CNN or 3D CNN is limited by the number of frames in a single input sequence, which is often subject to memory constraints. Longer sequences need to be partitioned into shorter segments to accommodate the CNN input size. The CNN will not form any connection between different input segments, which means that it can only detect local temporal features. LSTM, on the other hand, can handle arbitrary length inputs. With the help of internal hidden states, LSTM can recall information obtained from previous inputs for a long time, making it more suited for the task of velocity estimation.

We also demonstrated that the Deep-SMV approach substantially improves super-resolution reconstruction speed, particularly for regions with high vessel densities, allowing for real-time visualization of the microvascular flow dynamics. This is a particularly enticing accomplishment for super-resolution ultrasound imaging, potentially enabling the translation of the technology into functional ultrasound applications where high temporal resolution is critical for monitoring vascular response(s). Although other DL-based approaches have demonstrated rapid localization times, these solutions still rely on conventional ULM pairing and tracking algorithms to estimate velocity. The processing time burden of these algorithms scale with the MB count, making these solutions ineligible for dynamic real-time velocimetry applications. The Deep-SMV network processing time does not depend on the MB concentration, performing consistently across multiple imaging scenarios. This is evident in the performance gain of Deep-SMV in comparison to ULM for the imaging experiments presented in this manuscript: Deep-SMV modestly reduced processing time for the flow channel phantom (a low vessel density application) and yielded the best improvement for the *in vivo* cases, where the processing time requirement was reduced by an order of magnitude.

Although the Deep-SMV approach shows substantial promise over conventional ULM and competing DL solutions, the results presented in this manuscript must be understood within the context of the limitations of the study. The most critical is that there is no established gold-standard technique for measuring microvascular flow velocity to validate the output of the Deep-SMV network. As a compromise, we have used the velocity results of the much more computationally expensive ULM reconstructions as a reference standard; however, this approach comes with some caveats. As discussed, ULM processing is susceptible to spurious MB localizations which may bias the velocity point estimates. We have attempted to account for this by averaging the ULM velocity references across a long acquisition time, which should reduce the impact of this source of error. The network was trained using Field II simulations of MB flow, where the velocity distribution was taken from experimental ULM reconstructions of vasculature. This implicitly tunes some aspects of the Deep-SMV network to the parameters used in the initial ULM reconstruction and is also dependent on the assumptions applied during vascular modeling (e.g., laminar flow). This is demonstrated directly by the gain in Deep-SMV performance for organ-specific simulation data in comparison to more generic vascular flow training (**Figure 4**). Field II also cannot simulate nonlinear MB responses. Furthermore, although Deep-SMV demonstrates better handling of high MB concentrations, it is still susceptible to signal interference of densely packed MBs leading to a retrograde flow aliasing artifact in some larger vessels. The incorrect flow direction is also apparent in ULM reconstructions, but the long acquisition duration can reduce this effect by accumulating data from the wash-out phase of contrast enhancement. We are currently exploring adjustments to the Deep-SMV output to better encode for the circular flow angle topology to help address this limitation.

In the current training setup, we used SSIM loss, which is designed to provide a similarity measurement between two images that reflects how humans perceive them. Although SSIM has been successfully applied to various image processing tasks, it is not necessarily the most efficient in capturing errors in our desired output data. We believe that a loss function specifically tailored for velocimetry can potentially improve the performance of Deep-SMV.

Deep-SMV provides efficient, effective, and fast super-resolution microvascular structural and velocity reconstruction without requiring MB localization, pairing, and tracking. By eliminating this inefficient ULM processing pipeline, Deep-SMV can capitalize on the rich spatiotemporal information present in the input IQ data, enabling real-time super-resolution imaging that is robust to a wide range of imaging conditions. This technique demonstrated a high temporal resolution, permitting the application of super-resolution velocimetry to functional imaging scenarios and enabling super-resolved pulsatility mapping. By relaxing the strict imaging paradigm that ULM typically requires, Deep-SMV can facilitate the translation of super-resolution ultrasound to a wider range of experimental and clinical applications.

## Author contributions

XC, MRL, and PS designed and wrote the paper. NCS and DL prepared the mouse model and performed craniotomies. XC, MRL, ZD, and PS performed the ultrasound imaging on mice. CH and SC designed the mouse kidney study and protocol, prepared the mouse kidney model, and performed the ultrasound imaging. CH and SC evaluated and preprocessed the mouse kidney ultrasound data. TMF designed and produced the mouse tumor model used in this study. ZD designed the ultrasound transducer holder and programmed the motorized imaging stage. XC, MRL, and PS developed and applied the super-resolution ULM algorithm.

## Data Availability

The data that support the findings of this study are available from the corresponding authors on request.

## Competing Interests

The authors declare no competing interests.

## Funding

The study was partially supported by the National Cancer Institute, the National Institute of Biomedical Imaging and Bioengineering, the National Institute on Deafness and Other Communication Disorders, and the National Institute on Aging of the National Institutes of Health under grant numbers R00CA214523, R21EB030072, R21DC019473 and R03AG059103, as well as a grant to DL from the Kiwanis Neuroscience Research Foundation, and by the Seed Grant from the Cancer Center at Illinois. The content is solely the responsibility of the authors and does not necessarily represent the official views of the National Institutes of Health. NCS and MRL are supported by Beckman Institute Postdoctoral Fellowships.

## Methods

### Deep-SMV architecture and design

Our NN model adopts the convolutional LSTM-UNet structure, where the bottleneck layers of the classical UNet were replaced by LSTM layers (**Figure 1, Supplementary Figure 1**). The feature extraction path of the network starts with an input block that contains two convolution layers with 3*3 kernel size, each followed by a batch normalization layer and a Rectified Linear Unit (ReLU) activation function. The encoder block contains a 2*2 max pooling layer, two 3*3 convolution layers each followed by a batch normalization layer and ReLU activation function. Each encoder block reduces all spatial dimensions of its input by a factor of two and expands the feature dimension by a factor of two. Each frame of 2-D spatial input will be fed into the feature extraction blocks separately. The resulting feature maps were concatenated along the temporal dimension before being fed to the convolutional-LSTM layers as a feature map sequence.

The convolutional-LSTM block takes a new input and the previous hidden state output as it inputs, while maintaining an internal cell status variable. The inputs will first go through a 3*3 convolution layer. The convolution layer output will be split along the channel dimension into four parts, where three will go through a sigmoid activation function and operate as the forget gate, the input gate, and the output gate, respectively. A hyperbolic tangent (tanh) function will be applied to the remaining part. The forget gate determines which part of the previous cell status will be discarded. The input gate determines whether parts of the new input contribute to updating the cell status. A tanh activation function will be applied to the new cell status. Finally, the output gate determines how the cell status propagates to the output. The convolutional LSTM block will reduce the input sequence along the temporal dimension.

The decoder block is constructed similarly to the encoder block, with the max pooling layer replaced by spatial upsampling layer. Each decoder block will expand the spatial dimension of its input by 2 and reduce the channel dimension by 2. The output block is a 3*3 convolution layer with output channel size of 2. The final output of the network is a predicted blood flow velocity map with the same spatial dimension as the input data.

We used a combined mean squared error (MSE) and structural similarity (SSIM) as the loss function for training the NN, defined as:

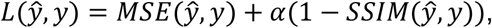

*ŷ* denotes the predicted velocity map and y denotes the ground truth velocity map. The MSE loss is the squared L2 distance between the prediction and ground truth averaged over all pixels:

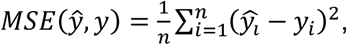

where the subscript i denotes the value of the ith pixel in the corresponding map. The SSIM is defined as

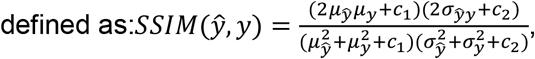

where *μ*_*ŷ*_ *μ*_*y*_ are the mean pixel values of the images, *σŷ σ*_*y*_ are the standard deviations, and *σ*_*yy*_ is the covariance of *y* and y. *c*_1_and *c*_2_ are constants to enforce numerical stability. We use (1-SSIM) in the loss function so that the structural similarity will be maximized in training. The SSIM scaling factor α is set to be 0.1.

### CAM optical and mouse brain ULM image-based training data simulation

The training dataset of our NN model was simulated using CAM optical images, as well as conventional mouse brain ULM results to serve as a vessel structure template.

#### CAM optical image acquisition

Optical imaging was performed using a Nikon SMZ800N stereomicroscope (Nikon, Tokyo, Japan) with a Plan Apo 1x objective at 1x zoom. Images were captured at 16 bits using an attached DS-Fi3 microscope camera, digitized using NIS Elements D software, and saved as .nd2 format. These were later exported as RGB .tiff files for external analysis (**Supplementary Figure 2a**). The optical imaging field of view was 11.01 mm x 15.48 mm yielding 2048-by-2880 pixel images.

#### CAM Vessel flow dynamics model

A binary vessel map of each CAM vasculature (**Supplementary Figure 2a**) was segmented by applying the *threshold_adaptive* function available in the scikit-image python package (29) to the green channel of each acquired optical image (**Supplementary Figure 2a**) as it provides the best contrast for blood vessels. Non-overlapping regions of 256-by-256 pixels were sampled from 35 2048-by-2880 pixel binarized CAM optical images, resulting in 3080 binarized ROIs, from which 1356 regions with acceptable segmentation performance were manually selected for further processing.

Each binary vessel map was skeletonized using the medial axis *skeletonize* function, also available in the scikit-image package. The medial axis *skeletonize* function returns the skeletonized image (**Supplementary Figure 2c**), as well as the distance transform map (**Supplementary Figure 2d**) which can be used as an estimation of local width of the vasculature. The skeleton image is converted to an undirected graph G = {V, E}, where E is the set of edges that represents vessel segments, and V is the set of vertices that represents junctions of the vessel segments (**Supplementary Figure 2e**). The graph is stored as a NetworkX (30) undirected graph. Conversion from skeleton image to NetworkX graph is achieved using the *sknw* function available in the ImagePy image processing framework (31).

The undirected vessel graph was then separated into a set of subgraphs, where each subgraph is a connected component of G. A spanning tree (a connected acyclic subgraph that covers all vertices in the original graph) was computed for each subgraph using the breadth-first search (BFS) algorithm starting from a source point defined as the degree-1 vertex with the largest local width in the corresponding distance map from the skeletonization step. Directions of the edges in the tree were assigned to start from the vertex that appeared earlier in the BFS traversal and point to the vertex that appeared later in the BFS traversal. The final directional vessel graph (**Supplementary Figure 2f**) is a forest that contains all the directional spanning trees obtained from each connected component of the undirected vessel graph.

#### Calculating reference velocity

We used the CAM ULM result as a reference to calculate a distribution of blood flow velocity on the central line of vessels for a given radius. We apply adaptive thresholding on the ULM velocity map to generate a binary vessel map, then perform medial axis skeletonization to obtain a local vessel radius estimation for each pixel on the medial axis of the vessel structure. All local radius-velocity pairs of pixels on the medial axis were sorted in ascending order of the local distances. The estimated reference velocity on the center line of a vessel with radius d was drawn from a normal distribution N(µ, σ^2^), where µ is the average velocity of all radius-velocity pairs with radius within d±0.5 pixels range in the list of radius-velocity pairs, and σ^2^ is the variance of the selected pairs.

#### Mouse brain ULM-based flow dynamics model

To study the effect of tissue-specific vascular structures in the training dataset, in addition to CAM-based vessel flow model, we also generated a simulation training set using mouse brain ULM images. 40 mouse brain ULM images (imaging field of view 7.17mm axial by 5.96mm lateral; λ/10 interpolation) were sampled as 256*256 pixels regions with a step-size of 128 pixels. The ULM images were generated from accumulated tracks of individual MBs, therefore will contain gaps and fragments even after long accumulation. For each sub-region, we apply *remove_small_holes* function from the skimage.morphology python package to fill the gaps and holes in ULM microvessel density map, followed by *remove_small_object* to clean up the remaining fragmented tracks. We then apply the same binarization-skeletonization procedure as described in the CAM-based simulation case and convert the cleaned-up vessel images into undirected graphs.

The undirected graphs were converted into directed vessel flow dynamics model with the same fashion as discussed in the CAM-based simulation case, except for velocity assignment. Here, we can directly use the corresponding mouse brain ULM velocity map. For each vessel segment, we calculate the average velocity of all the non-zero pixels in the ULM velocity map within the area of the segment and use it as the velocity along the center line of the vessel segment in the directed vessel flow graph.

#### Simulating MB motion

Once the vessel flow dynamics model was constructed, we can use it to generate a sequence of MB locations that travel in the vessel network. Each MB was stored as a tuple (e_orig, e_curr, d_ax, d_lat, amp), where e_orig is the vessel segment (edge) that it originated from, e_curr is the edge that it is currently traveling along, d_ax is the axial distance with respect to the current edge, d_lat is the lateral distance with respect to the current edge, and amp is the amplitude of the MB. The spatial location of the MB with respect to the entire field-of-view (FOV) can be easily inferred from e, d_ax and d_lat. The MB location was updated using the following rules:

- Initialization: place 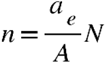 MBs on each edge, where *a*_*e*_ is the volume of the current vessel segment, estimated using the local radius at points on the edge e, 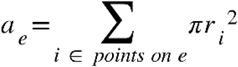. Set both e_orig and e_curr to be e. d_ax and d_lat were both randomly chosen so that the MB stays within the current vessel segment. Amp was randomly selected from a uniform distribution on (0, 1]. E_orig and amp will remain unchanged until the MB is removed from the vessel network.
- For each timestep, perform the following for each MBs:
  - Calculate the axial velocity along the direction of the current vessel segment of each MB according to the Poiseuille flow condition, 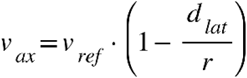, where *v*_*ref*_ is the reference flow velocity on the centerline calculated using the method described in the previous section, and r is the local radius of the vessel segment. Calculate the displacement *v*_*ax*_·Δ*t* where ;Δ*t* is the temporal resolution.
  - If the displacement is smaller than the remaining length of the current edge:
    ▪ Update d_ax with the displacement.
    ▪ Update d_lat with a number randomly generated small distance so that the MB still stays within the vessel segment.
    ▪ E_curr remains unchanged.
  - If the displacement is larger than the remaining length of the current edge:
    ▪ If the current edge ends at a leaf vertex (i.e., no outgoing edge from the vertex), add a new MB to vessel segment e_orig of the current MB, remove the old MB from the vessel network.
    ▪ If the current edge has outgoing adjacent edges, the MB will be moved to one of the adjacent edge *e* _*aj*_ with probability, 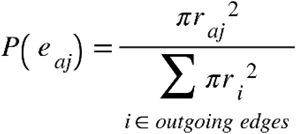 where r is the local radius of the starting vertex of each edge. Subtract the remaining length on the current edge from the total displacement. Update e_curr to e_aj. Repeat the steps above until the remaining displacement is less than the remaining length of the current edge. Then, update d_ax and d_lat with the same method in the previous step using the final remaining displacement.
  - Calculate the spatial coordinates of each updated MB using e, d_ax and d_lat. Store the list of locations and amplitudes of each MB. After repeating the update steps for a desired duration of time, save the list of MB locations and amplitudes of each timestep as input to Field-II simulation.

#### Field-II simulation

Ultrasound simulation of MB motion was performed in Field-II (23, 24) using the sequence of MB locations and amplitudes obtained in the previous simulated MB motion step. For each timestep, we generate one frame of ultrasound simulation of point scatterers specified by the corresponding list of locations and amplitudes in the sequence. The imaging sequence was configured according to the CAM ultrasound imaging experiment setting, with a high frequency linear array transducer, 20MHz center frequency, and 9-angle plane wave compounding. Detailed simulation parameters were summarized in **Table I**.

**Table 1.**
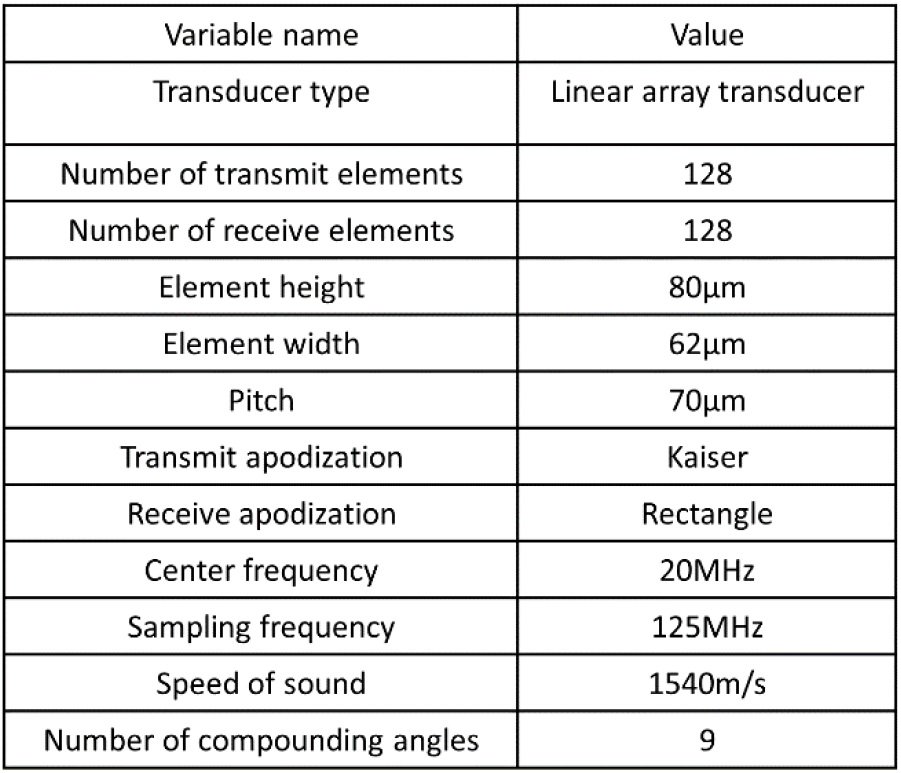
Field-II Simulation Parameters

#### Deep-SMV training

Training samples of 16 frames in the temporal dimension, and 256 * 256 pixels in the spatial dimension were fed to the model in mini-batches of 4 samples per batch. The model was trained with an Adam optimizer to minimize the SSIM loss between the model output and the ground truth using back-propagation with a learning rate of 0.001. The ground truth contains two channels of the magnitudes in pixels/ms and angles of the flow velocity map. To avoid complications of training with negative values in the ground truth, the angle map which originally contains values in the range [−*π, π*] was scaled and shifted to the scale [0, 1]. The model was trained for a total of 100 epochs.

### Ethics approval

All procedures performed on mice at the University of Illinois Urbana-Champaign in this manuscript were approved by the Institutional Animal Care and Use Committee (IACUC). All experiments were performed in accordance with the IACUC guidelines. Mice were housed in an animal care facility approved by the American Association for Assessment and Accreditation of Laboratory Animal Care.

All experimental procedures involving animals at the Mayo Clinic were performed according to the Guide for the Care and Use of Laboratory Animals (NIH publication Nos. 80–23, revised 1996) and followed the institutional ethical guidelines for animal experiments. All experiments were approved by local Ethics Committee of the Mayo Clinic.

No IACUC approval was necessary to conduct the chicken embryo imaging presented in this manuscript, as avian embryos are not considered to be “live vertebrate animals” under NIH PHS policy.

### Ultrasound imaging

All flow phantom, chicken embryo, and mouse brain ultrasound data were acquired with a Vantage 256 system (Verasonics Inc., Kirkland, WA) with a L35-16vX high-frequency linear array transducer. Imaging was performed with a center frequency of 20 MHz, using 9-angle plane wave compounding (1-degree increments) with a post-compounding frame rate of 1,000 Hz. Ultrasound data was saved as in-phase quadrature (IQ) datasets of 1,600 frames each for external processing. Mouse kidney imaging was performed at a center frequency of 25 MHz, and a post-compounding frame rate of 500 Hz.

### Microbubble flow phantom

A pair of holes were drilled on the opposite walls of a clear rectangular plastic container. Stainless steel dispensing needles were attached to the outside wall of the container on top of the holes using epoxy adhesive. A stainless-steel rod with a diameter of 500μm was inserted horizontally into the container through the needle openings. A 20% gelatin mixture was poured into the container until the surface was approximately 5mm above the metal tube. The phantom was placed in a 4ºC refrigerator until the gelatin completely solidified. The metal tube was removed prior to the experiment to create a flow channel, the diameter of which constricted to approximately 450μm during the gelatin solidification process. The inlet to the flow channel was connected to a programmable syringe pump (NE-300, New Era Pump Systems Inc., Farmingdale, NY) with a soft plastic tube to provide constant flow volume rate through the channel. A clinically available ultrasound contrast agent (DEFINITY®, Lantheus Medical Imaging, Inc.) was diluted 1000-fold with 0.9% saline (0.9 sodium chloride, BD, Franklin Lakes, NJ) and was perfused through the flow channel. Volume rates of 40-180μL/min with 20μL increment was used in this experiment. The L35-16vX transducer was placed at the top surface of the phantom and positioned to provide a longitudinal view of the flow channel (**Supplementary Figure 3a**). For each volume rate, we waited until a stable flow was formed before starting the acquisition of 1.6s duration (1,600 frames).

### Ex ovo CAM preparation

Fertilized leghorn chicken eggs (*Gallus gallus domesticus*) were provided locally by the University of Illinois Poultry Research Farm. Eggs were housed in a tilting incubator (Digital Sportsman Cabinet Incubator 1502, GQF manufacturing Inc.) until embryonic development day 4 (EDD-04). The eggshells were removed using a rotary Dremel tool and the egg contents (yolk, albumin, and embryos) were transferred into plastic weigh boats. The now *ex ovo* chicken embryos were then placed into a humidified incubator (Darwin Chambers HH09-DA) until imaging on EDD-17.

A glass capillary needle was produced using a PC-100 glass puller (Narishige, Setagaya City, Japan) and borosilicate tube (B120-69-10, Sutter Intruments, Novato, CA, USA) to aid contrast MB injection. The glass needle was attached to a 1mL syringe using Tygon R-3603 laboratory tubing, and the surface vasculature of the CAM was injected with 70 μL of DEFINITY® at clinical bolus concentration immediately prior to ultrasound imaging. The L35-16vX transducer was placed on the side of the plastic weigh boat to image the surface vasculature of the CAM (**Figure 2g**).

### Mouse kidney imaging

Kidney imaging was performed on an anesthetized mouse (2% isoflurane mixed with oxygen) following a 20 μL bolus injection of Bracco Lumason (Bracco, Milan, Italy) microbubble solution via tail vein. The L35-16vX was positioned to produce a long axis view (**Figure 6a**). A total of 12 IQ datasets each with 720 frames of data (8,640 frames in total) corresponding to a data length of 17.28 seconds, were stored for post-processing. Contrast power images were generated by accumulating the MB signal along the temporal dimensions, and color Doppler images were generated using the technique by Loupas *et al*. (32).

### Mouse tumor preparation and imaging

Athymic nude mice were inoculated with 5 × 10^6^ Hepa1-6 cells subcutaneously into their right flank. After 10 days of growth, the mice were anesthetized with isoflurane and were then secured to a heated imaging platform, and a tail vein catheter was inserted to aid with MB delivery. The L35-16vx transducer was placed to produce the largest cross-section of the tumor (**Figure 6e**), and then a 50uL bolus of DEFINITY® was injected before imaging. IQ data was stored as 1,600 frame acquisitions for post-processing. Contrast power and color Doppler images were produced as above.

### Mouse craniotomy preparation

Anesthesia was induced by placing each CBA/CAJ (stock # 000654) mouse in a gas induction chamber supplied with 4% isoflurane mixed with medical oxygen. Then the mice were placed in a stereotaxic frame with nose cone supplying 1-2% isoflurane with oxygen for maintenance. Homoeothermic heating pads were used to maintain body temperature at 36.5°C. The surgical area was shaved and cleaned with iodine before being injected with Lidocaine (1%) for local anesthesia. Ear bars were used to secure the mouse head to the stereotaxic imaging stage. The scalp of the mouse was removed, and a cranial window was opened on the left side of the skull using a rotary Dremel tool, starting at the sagittal suture and moving laterally to expose the cerebral cortex. The tail vein of the mouse was cannulated with a 30-gauge catheter and vessel patency was confirmed with a 0.1 mL injection of sterile saline. The L35-16vX transducer was positioned to acquire a coronel imaging plane of the brain through the cranial window (**Figure 4, Figure 5**). Fresh DEFINITY® was perfused via the tail vein catheter at a rate of 10 µL/min using a programmable syringe pump (NE-300, New Era Pump Systems Inc., Farmingdale, NY). The MB solution was mixed every 5 minutes using a magnetic stirrer to maintain a constant MB concentration during the experiment. A total of 40 acquisitions (64,000 frames, or 64 seconds of data) were acquired for this imaging plane. Electrocardiogram (ECG) signal was acquired using an iWorx (Dover, NH, USA) IA-100B single channel biopotential amplifier with C-MXLR-PN3 platinum needle electrodes inserted into the legs of the animal.

### Conventional ULM image processing and analysis

MBs were extracted for each ultrasound IQ dataset by applying a spatiotemporal singular value decomposition (SVD)-based clutter filter (25, 33–35), where the low-order singular value threshold representing tissue signal was determined adaptively (36). A noise-equalization profile was applied to account for depth-dependent effects (37). An isolated MB signal was manually identified and fit to a multivariate Gaussian function to represent the point-spread function (PSF) of the system. A MB separation filter (25) was applied to each SVD-filtered dataset, and then the imaging data was spatially interpolated to an isotropic 4.9 µm axial/lateral resolution using spline interpolation (38). MBs were then localized by applying a 2D normalized cross-correlation between the empirical PSF and each frame of interpolated IQ data. MB centroid pairing and frame-to-frame trajectory estimation was performed using the uTrack algorithm (39), or a Kalman filtering algorithm (27), where velocity was calculated from centroid displacement. Each processed data acquisition was accumulated into a final reconstruction.

## Notes

### Competing Interest Statement

The authors have declared no competing interest.

