## Supplemental Figures for "Localization free super-resolution microbubble velocimetry using a long short-term memory neural network"

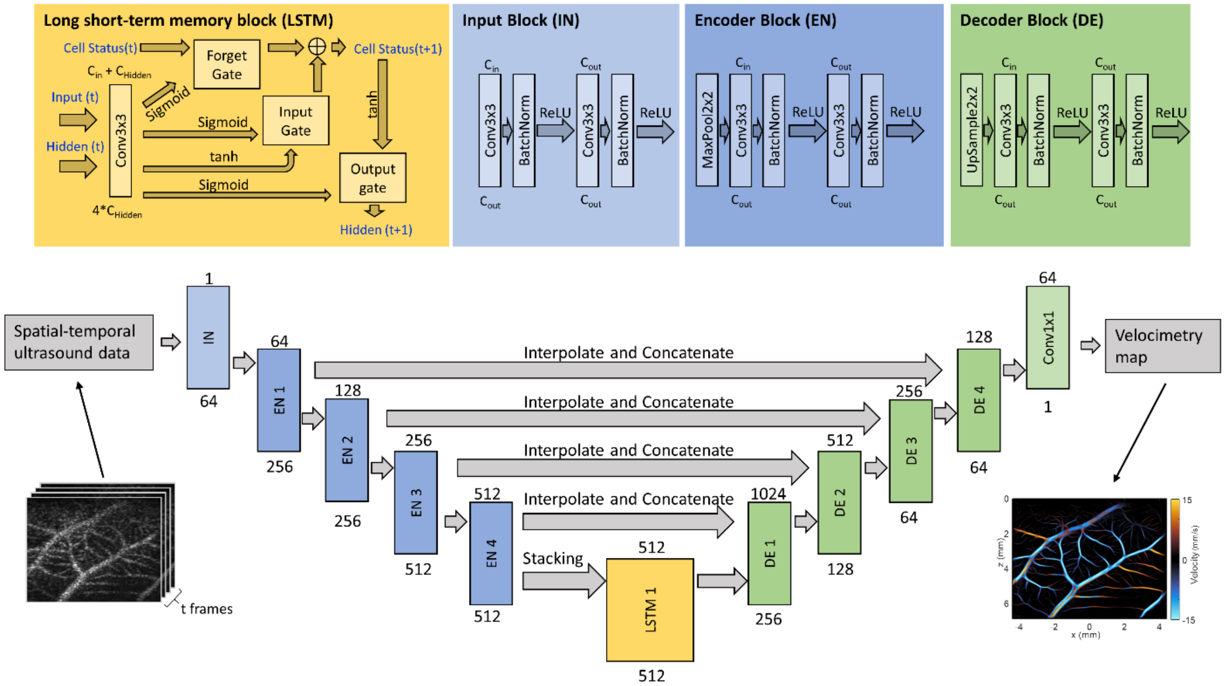


**Supplementary Figure 1. Deep neural network architecture for Deep-SMV.** The main network design is a classic UNet structure with long short-term memory (LSTM) blocks in the bottleneck layers to provide flow velocity measurements, with input consisting of spatial-temporal (2D spatial + time) ultrasound data and output of super-resolution velocity and structure maps. The LSTM, input, encoder, and decoder block designs are provided in detail at the top of the figure. The text above each block indicates the input channel size to the unit, and the text under each block indicates the output channel size from the unit.

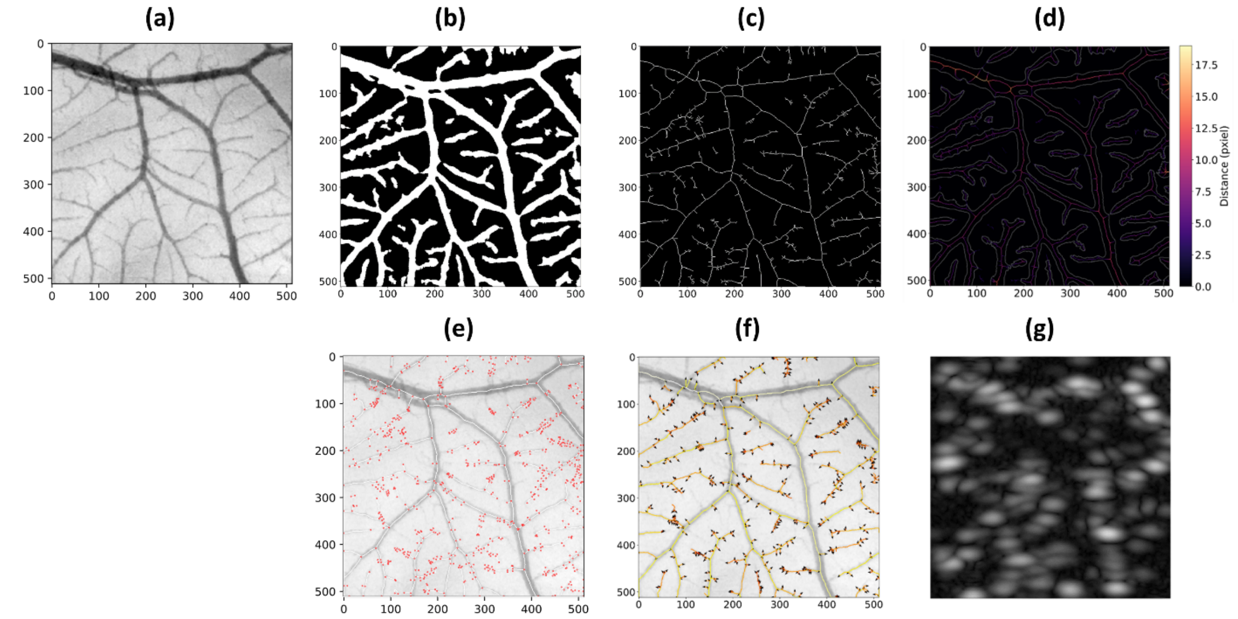


**Supplementary Figure 2. CAM-based MB flow simulation procedure. (a)** Grayscale image of the green channel pixel intensity of a 512-by-512 pixels region in a CAM optical image. **(b)** Binary segmentation result of (a) by applying adaptive thresholding. **(c)** Skeleton of (b) obtained by applying medial axis skeletonization on (b). **(d)** Distance transform of (b) obtained by applying medial axis skeletonization on (b). **(e)** Undirected graph constructed from the skeleton (b), displayed on top of the original CAM image. Lines represent edges and dots represent vertices. **(f)** Final directed vessel graph with arrows indicating flow direction on each vessel segment. **(g)** Example frame of Field-II ultrasound simulation from MB flow simulation.

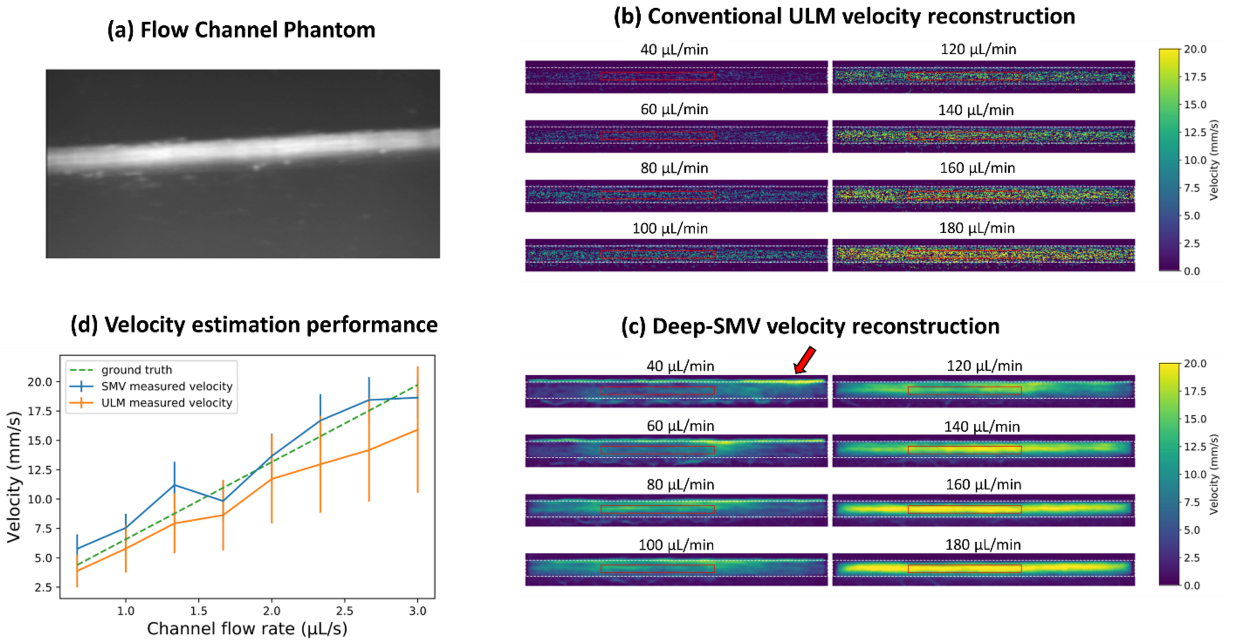


**Supplementary Figure 3. Flow channel phantom validation. (a)** A contrast power Doppler image of the experimental flow phantom. **(b)** Conventional ULM localization and tracking was able to sparsely populate the flow channel with velocity estimates. Due to the inefficient use of MB signals, a much longer acquisition time would be required to completely scout out the channel lumen. **(c)** Deep-SMV was also able to provide velocity estimates for this flow channel and was much more efficient in using the MB signal, resulting in a higher proportion of reconstructed luminal space. **(d)** The performance of velocity estimates for different flow volume rates between ULM and Deep-SMV, in comparison to the theoretical gold-standard. ULM demonstrated an underestimation bias for velocity estimates, particularly for high flow volume rates. The analysis ROIs were placed to avoid the upper edge enhancement artifact from buoyant MBs at low flow volume rates (arrow).

**Supplementary Video 1. Deep-SMV CAM vessel pulsatility video.** This video demonstrates the super-resolution functional vascular imaging capability of Deep-SMV processing. During the time course of ultrasound acquisition (1.6 seconds) a cyclic pulsatility is visible across the CAM vessel network at a frequency approximately matching the expected cardiac cycle rate at this stage of embryonic development (2-3 Hz).
